# Functional asymmetry in the central brain regions in boys with Attention Deficit Hyperactivity Disorder detected by Event Related Potentials during performance of the Attentional Network Test

**DOI:** 10.1101/118380

**Authors:** Dimitri Marques Abramov, Carlos Alberto Mourão, Carla Quero Cunha, Monique Castro Pontes, Paulo Ricardo Galhanone, Leonardo C. deAzevedo, Vladimir V. Lazarev

**Author notes:** Corresponding Author: Dimitri Marques Abramov, Insituto Nacional da Saúde da Mulher, da Criança e do Adolescente Fernandes, Figueira, Fundação Oswaldo Cruz, Av. Ruy Barbosa, 716, Flamengo, Rio de Janeiro, RJ, Postal Code: 22.250-020.

## Abstract

**Background:** Various functional asymmetries detected by different neurophysiological and neuroimaging methods have been reported in the literature on the Attention Deficit Hyperactivity Disorder (ADHD), some of them pointing to the right hemisphere activity. In our attempt to discriminate the ADHD patients from normal subjects by hierarchical clustering of behavioural, psychological and event related potential (ERP) variables, the late P3 component of potentials from the right central region (C4) proved to be one of the most informative parameters (in preparation for publication). Here, we have studied the differences in ERPs between the left (C3) and right (C4) central leads and relation of this asymmetry to ADHD diagnosed using DSM.

**Methods:** 20 typically developing (TD) boys and 19 boys diagnosed with ADHD according to DSM-IV-TR, aged 10-13 years, were examined by the Attentional Network Test (ANT), with simultaneous recording of the respective ERPs. The intergroup differences in the ERP amplitude parameters in the left (C3) and right (C4) central channels and in the difference in these parameters between the two channels (C3 minus C4) were accessed. These characteristics were compared to the subjects DSM scores and ANT performance.

**Results:** The target-related potentials late characteristics from the C4 and C3 did not shown significant difference between the groups. The difference between ERPs of the C3 and C4 channels inside the interval of 40-290 ms after target onset was larger in the ADHD group than in control, mainly for incongruent target condition. This asymmetry and right late component were correlated with DSM scores, mainly to hyperactive and impulsive criteria.

**Conclusion:** In ADHD patients, the results suggest ERP pattern of right-side functional predominance in the motor control, which correlates to DSM scores, mainly to hyperactive and impulsive criteria.

## 1. Introduction

The cerebral mechanisms of the Attention Deficit Hyperactivity Disorder (ADHD) have been related to the alterations in the frontal executive control upon sensory-motor functions (1, 2, 3). Some evidences of asymmetric character of these alterations were found in the right frontal lobe, both in the prefrontal cortex (4) and deeper, in the caudate nucleus (5), where the neuroimaging data correlated with the Diagnostic and Statistical Manual of Mental Disorders (DSM) scores. The cortico-caudate circuits are largely involved in the behavior and motor control (6, 7). ADHD and hiperkinetic disorders, (8), as well as other movement disorders (9) correlate with cortico-striatal alterations. A striking feature of ADHD patients is their behavioral hyperactivity and impulsivity (1,8), which serve criteria for ADHD diagnosis in the DSM (10, 11).

In our previous EEG research, we also observed the asymmetrical differences between ADHD and control groups, with the signs of relative inactivation of the left fronto-temporal cortex known as responsible for the voluntary attention (Lazarev et al., 2016). This left-side ‘inactivation’ could be compensated by relatively higher activation of the contralateral cortex. This can partially explain the leading role of the ERP data from the right fronto-temporal and particularly right central motor regions in discriminating the ADHD patients from the control subjects by hierarchical clustering of behavioral, psychological and ERP variables observed in our other research (in preparation for publication).

Here, we have studied the scalp ERPs from the left and right central brain areas during performance of the Attentional Network Test (ANT), a forced two-choice test addressed to multiple dimensions of attention (vigilance, spatial orientation and conflict resolution) according to Posner’s theory of Attentional Networks (12–17). In the ANT, motor performance, evidenced by reaction time (RT), is manipulated by the information contained in the cue and target stimuli (12). The ANT-related ERPs proved to be sensitive to ADHD (16).

The objective of this paper is a preliminary report about functional asymmetry in the central motor areas related to ADHD patients’ behavior during the ANT performance. This is a partial presentation of the results, which are in preparation for publication and include ERPs data recorded from various cerebral areas.

## 2. Methodology

### 2.1. Subjects

Thirty two boys, aged 10-13 years, were sampled according to DSM-IV-TR: 19 with ADHD and 20 typically developing (TD) subjects. All them were free of psychotropic medicines for the last 30 days, without history of neither chronic diseases nor psychiatric disorders, as screened by K-SADS-PL (18). Their estimated intelligence quocient (I.Q.) was > 80 (see below).

The study was approved by the Ethics Committee of the National Institute of Women, Children and Adolescents Health Fernandes Figueira. All the primary caregivers gave written informed consent after receiving a complete description of the study. The boys also gave their oral assent.

### 2.2. Clinical and psychological examination

Each subject was evaluated by a structured interview where their caregivers were shown the DSM-IV-TR criteria, and were instructed to point out carefully whether or not each specific criterion was an exact characteristic of their children’s behavior. If there was any doubt or hesitation about any item, it was disregarded. Thus, subjects were classified in accordance with the DSM-IV-TR.

The I.Q. was estimated by Block Design and Vocabulary, subtests from the Wechsler Intelligence Scale for Children, 3^rd^ version (WISC-III) (19, 20). In the previous study, the I.Q. scores estimated by these two subtests showed very high correlation with the results from the full WISC scale (20).

### 2.3. Experimental procedures

The ANT version adapted for children, with little fishes instead of traditional arrows, in line to Kratz et al. (15), were used (15b). The ANT is a forced two-choice test where the subject is instructed to look fixedly at the central fixation point and observe the horizontal orientation of a target stimulus flanked by distractors (two similar fishes at each side, all with the same orientation) and preceded or not by a cue signal, which informs where and/or when the target appears. The horizontal orientation of the target to the right or to the left was equiprobable and the same (congruent) or opposite (incongruent) to the orientation of distractors. There were three equiprobable cue conditions corresponding to this signal’s position or its nonappearance: 1) at the subsequent upper or lower position of the target - Spatial cue condition; 2) at the central fixation point - Neutral cue condition; or 3) No-cue condition. The subject had to press promptly with his index or middle finger the left or right arrow key of the keyboard, according to the target horizontal orientation. The target appeared for 350 ms, 100 ms after the distractors. There was a random interval from 1 to 2 s between the trials. The time interval between the cue and target presentations was 1650 ms. The test was organized in 9 blocks, each block consisting of 24 trials, i.e. 8 trials of each cue condition, presented in random order. The first block was used only for training and the other eight were testing. More detailed information can be seen in (15b).

### 2.4. EEG acquisition

During the ANT performance, the subject’s EEG was recorded by a Nihon Kohden NK1200 EEG System at at paracentral (C3 and C4) sites according to the International 10/20 system, with monopolar reference to the linked earlobes (A1+A2). The impedance was below 10 kΩ. The EEG was recorded at a 1000 Hz sampling rate and resolution of 16 bits, with low-pass (0.5 Hz), high-pass (100 Hz) and notch (60 Hz) filters.

### 2.5. Data analysis

Here we have focused only on the C3 and C4 leads, located over motor areas, about 5 cm to the left and to the right from the vertex, respectively (21). The target ERPs (triggered at the target onset) and the arithmetic difference between the left (C3) and right (C4) waves were subject to analysis.

For the target ERPs, we estimated the maximum peak amplitude and calculated the sum of all positive and negative amplitudes inside the time window at 200 – 800 ms after the target onset (marked with bold black lines in Figure 1), which embraced the late ERP component. The latter parameter called ‘total amplitude’ (TA) was equivalent to the mean amplitude. We also considered the same measures for the asymmetry between the waves in C3 and C4 (‘C3 minus C4’ channel) inside the time window at 45 – 290 ms (Figure 2). The above ERP parameters were estimated for each cue condition and for all of them together, and also for both congruent and incongruent target position. We compared DSM scores, peak amplitudes, mean amplitudes, RTs and intraindividual variation of RT between the groups using the non-parametric Mann-Whitney U-test (alpha = 0.05), because no sample was regarded as normally distributed, once groups were too small (n=19 and 20).

**Figure 1.**
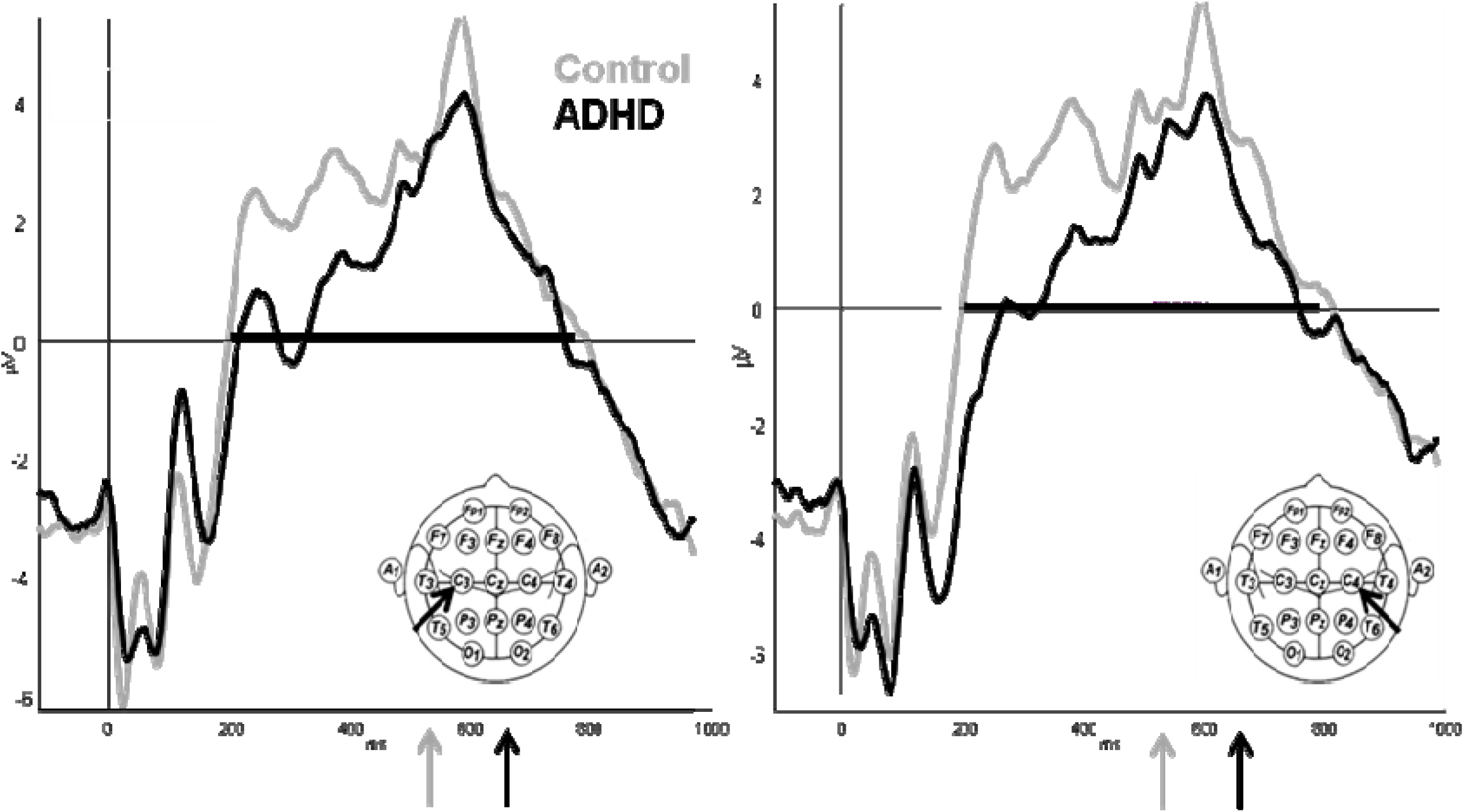
ERPs (μV) for left (C3) and right (C4) central channels (at left and right in figure) averaged across subjects of control (gray) and ADHD (black) groups. Bold black line on the abscissa shows time window with late ERP components. The mean RT for control and ADHD were marke d on graphs by arrows

**Figure 2.**
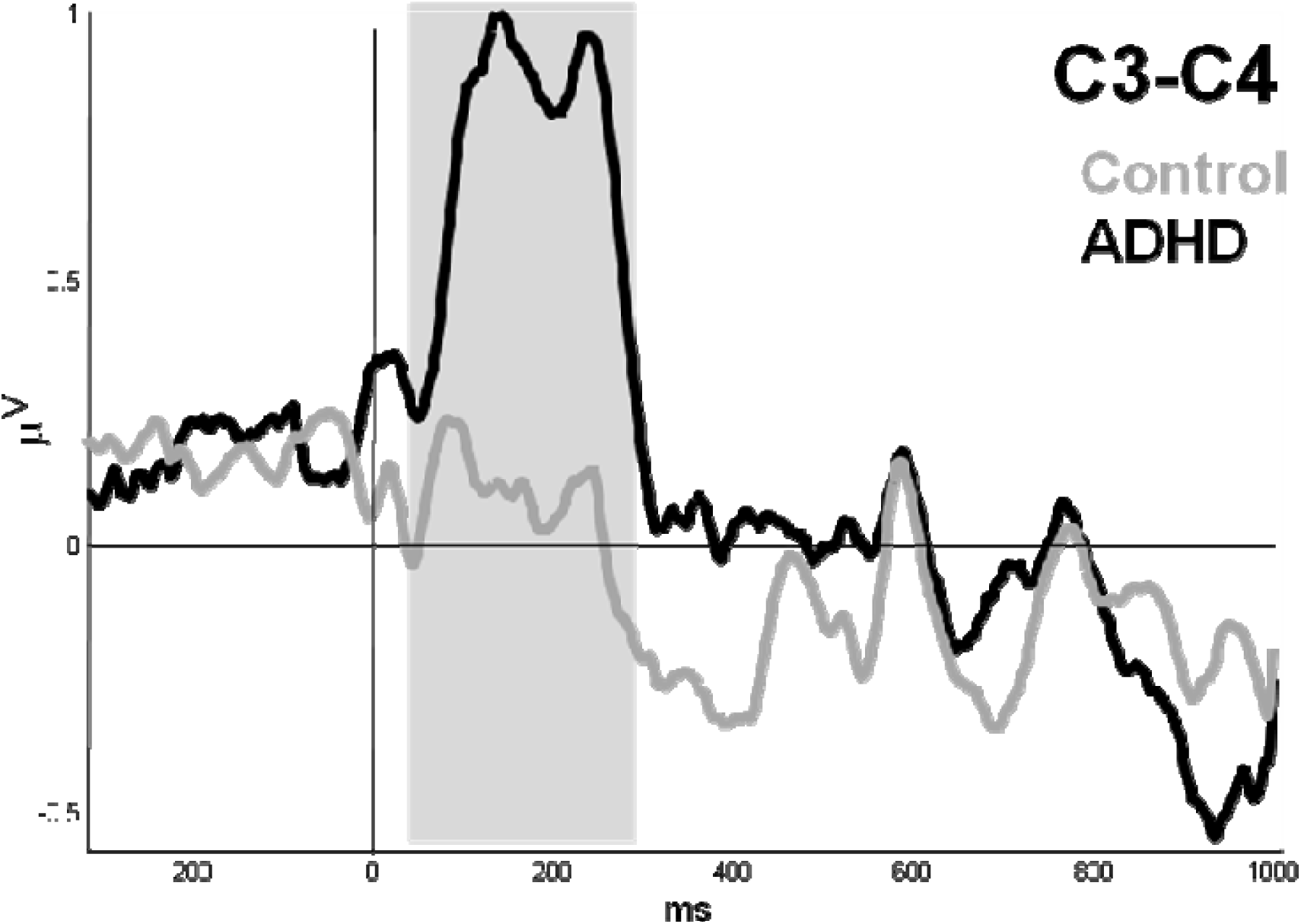
ERP asymmetry from ‘C3 minus C4’ channel, for control (gray) and ADHD (black) groups.The gray area shows the time window of int erest (45-290 ms).

The correlation coefficients and their probabilities, between the ERP characteristics and the mean RT were calculated using two-tailed Spearman’s Rank Test (ρ) for each group (n=19 and 20), and between ERPs and or DSM scores using two-tailed Pearson’s test (r) for all subjects (n = 39).

## 3. Results

The ADHD and TD groups have presented significant difference in the most of DSM scores, except for hyperactivity (p = 0.063) (Table 1).

**Table 1.** Difference between control and ADHD groups for DSM-scores

| DSM criteria | Control |  | ADHD |  | p-value |
| --- | --- | --- | --- | --- | --- |
|  | Mean | Std Dev | Mean | Std Dev |  |
| Innatentive | 2.40 | 1.66 | 7.21 | 1.27 | 0,000 |
| Hyperactivity | 1,70 | 1.17 | 2.73 | 2.02 | 0,121 |
| Impulsivity | 0.95 | 0.68 | 1.63 | 1.11 | 0,040 |
| Hyperactivity+Impulsivity | 2.60 | 1.63 | 4.36 | 2.90 | 0,051 |
| Total | 5.00 | 2.75 | 11.63 | 3.00 | 0,000 |

In the ANT, the average RT for all conditions was not significantly different (z-stat = 1.14; p = 0.255) between the control (0.56 ± 0.12 s) and ADHD (0.61 ± 0.14 s) groups. However, the intraindividual variation of reaction time was different (z-stat = 2.04; p = 0.042) between control (0.16 ± 0.06 s) and ADHD groups (0.23 ± 0.11 s)

The ERP waveforms for the ADHD and TD groups demonstrated early (from 20 to 200 ms) and large late (from 200 to 800 ms) target-related components that appeared bilaterally. However, in the ADHD patients, the late component achieved its maximum positive amplitude level about 330 ms later than in the controls being negative during initial 200 ms, although the maximum peaks in both groups had similar latency ∼ 600 ms (Figure 1). Among the amplitude characteristics of the late ERP component, there are no differences found.

The amplitude difference between the left and right ERPs (channel ‘C3 minus C4’) during the period from 45 to 290 ms after the target onset was larger in the ADHD than the control subjects (Table 2 and Figure 2). The peak amplitude and mean amplitude of this asymmetry was statistically different between groups for all cue conditions (p < 0.05), as well as for spatial, congruent and incongruent conditions separately (p < 0.05, table 2).

**Table 2.** Group averaged peak and total amplitude (sum of all amplitudes inside time window*, μV) of the target-related ERP asymmetry inside the time windows of interest and statistical significance of differences between the groups

|  | Control |  | ADHD |  | Z-stat | p |
| --- | --- | --- | --- | --- | --- | --- |
|  | Mean | Standard deviation | Mean | Standard deviation |  |  |
| All conditions – peak amplitude | -0,11 | 2,52 | 1,74 | 1,86 | 2,57 | 0,010 |
| All conditions – mean amplitude | 0,08 | 1,14 | 0,75 | 0,90 | 2,09 | 0,036 |
| No Cue – peak amplitude | -0,05 | 2,96 | 1,68 | 2,50 | 1,87 | 0,062 |
| No Cue – mean amplitude | 0,07 | 1,16 | 0,78 | 1,02 | 1,92 | 0,054 |
| Neutral cue – peak amplitude | -0,07 | 3,24 | 1,42 | 2,28 | 1,45 | 0,148 |
| Neutral cue – mean amplitude | 0,05 | 1,33 | 0,66 | 0,90 | 1,50 | 0,133 |
| Spatial cue – peak amplitude | 0,09 | 2,79 | 2,10 | 2,19 | 2,15 | 0,032 |
| Spatial cue – mean amplitude | 0,12 | 1,17 | 0,81 | 0,89 | 1,87 | 0,062 |
| Congruent target – peak amplitude | -0,27 | 2,77 | 1,85 | 1,97 | 2,26 | 0,024 |
| Congruent target – mean amplitude | 0,11 | 1,19 | 0,74 | 0,88 | 1,92 | 0,054 |
| Incongruent target – peak amplitude | -0,04 | 2,57 | 1,81 | 2,05 | 2,43 | 0,015 |
| Incongruent target – mean amplitude | 0,05 | 1,14 | 0,76 | 0,96 | 2,09 | 0,036 |

(\*) 45 - 290 ms after target onset for 'C3 minus C4'

Considering all subjects together, despite their groups, the asymmetry from the ‘C3 minus C4’ channel correlated with all DSM scores mainly for hyperactive and impulsive (table 3). The late component from C4 site was also correlated to hyperactive-impulsive criteria.

**Table 3.** Spearman’s and Pearson’s correlation analysis between DSM scores and event-related potentials (P3 at C3 and C4, and re spective asymmetry)

| DSM dimension | Event-Related Potential | Spearman's test |  |  |  | Pearson's test |  |  |
| --- | --- | --- | --- | --- | --- | --- | --- | --- |
|  |  | Control |  | ADHD |  | All subjects |  | Sig. |
|  |  | Rho | p(Rho) | Rho | p(Rho) | r | p(r) |  |
| Inattention | C3 - P3, peak ampl | 0,01 | 0,967 | 0,17 | 0,491 | -0,02 | 0,916 |  |
|  | C3 - P3, mean ampl | -0,27 | 0,258 | 0,20 | 0,410 | -0,17 | 0,312 |  |
|  | C4 - P3, peak ampl | -0,18 | 0,455 | 0,23 | 0,344 | -0,13 | 0,420 |  |
|  | C4 - P3, mean ampl | -0,30 | 0,204 | 0,25 | 0,301 | -0,21 | 0,191 |  |
|  | asymmetry, peak ampl | 0,45 | 0,045 | -0,41 | 0,083 | 0,45 | 0,004 | # |
|  | asymmetry, mean ampl | 0,37 | 0,112 | -0,12 | 0,629 | 0,35 | 0,028 | # |
| hyperactive + impulsive | C3 - P3, peak ampl | -0,03 | 0,884 | -0,09 | 0,714 | -0,17 | 0,307 |  |
|  | C3 - P3, mean ampl | -0,41 | 0,073 | -0,14 | 0,572 | -0,25 | 0,123 |  |
|  | C4 - P3, peak ampl | 0,01 | 0,961 | -0,28 | 0,249 | -0,33 | 0,040 |  |
|  | C4 - P3, mean ampl | -0,37 | 0,111 | -0,40 | 0,090 | -0,43 | 0,007 | ## |
|  | asymmetry, peak ampl | 0,40 | 0,082 | 0,39 | 0,098 | 0,46 | 0,004 | ## |
|  | asymmetry, mean ampl | 0,48 | 0,032 | 0,32 | 0,180 | 0,48 | 0,002 | ## |
| total score | C3 - P3, peak ampl | -0,02 | 0,937 | -0,09 | 0,721 | -0,11 | 0,492 |  |
|  | C3 - P3, mean ampl | -0,36 | 0,120 | -0,06 | 0,806 | -0,25 | 0,121 |  |
|  | C4 - P3, peak ampl | -0,10 | 0,678 | -0,23 | 0,351 | -0,26 | 0,104 |  |
|  | C4 - P3, mean ampl | -0,34 | 0,145 | -0,25 | 0,298 | -0,37 | 0,020 | # |
|  | asymmetry, peak ampl | 0,56 | 0,011 | 0,16 | 0,525 | 0,54 | 0,000 | ### |
|  | asymmetry, mean ampl | 0,55 | 0,012 | 0,21 | 0,385 | 0,49 | 0,001 | ## |

## 4. Discussion

The interhemispheric asymmetry was statistically different between the ADHD and control subjects showing the right lower amplitude in the ADHD. Moreover, the early asymmetry as right late component correlated with the DSM scores, mainly the hyperactivity+impulsivity ones. This points to a more intrinsic biological association with the clinical phenomenology.

